# Asgard ESCRT-III and VPS4 reveal evolutionary conserved chromatin binding properties of the ESCRT machinery

**DOI:** 10.1101/2021.11.29.470303

**Authors:** Dikla Nachmias, Nataly Melnikov, Alvah Zorea, Yasmin De-picchoto, Raz Zarivach, Itzhak Mizrahi, Natalie Elia

## Abstract

The ESCRT machinery drive membrane remodeling in numerous processes in eukaryotes. Genes encoding for ESCRT proteins have been identified in Asgard archaea, a newly discovered superphylum, currently recognized as the ancestor of all eukaryotes. This begs the question of the functional evolutionary origin of this machinery and its conservation across lineages. Here, we find that Asgard-ESCRT’s exhibit conserved DNA-binding properties, which is derived from recruitment of specific members. We show that Asgard-ESCRT-III/VPS4 homologs interact with one another inside mammalian cells, associate with chromatin, and recruit their counterparts to organize in discrete foci in the mammalian nucleus. This is congruent with human-ESCRT-III homologs. We find that human- and Asgard-ESCRT-IIIs associate with chromatin via the same N terminal domain, and that human-ESCRT-III can recruit Asgard-VPS4 to the nucleus to form foci. Therefore, ESCRTs possess chromatin binding properties that were preserved through the billion years of evolution that separate Asgard and human cells.

## INTRODUCTION

Encoded by all domains of life, the ESCRT machinery constitutes one of the most evolutionary conserved cellular membrane remodeling machines (*1*). ESCRT is a multiprotein complex that is composed of five subfamilies i.e. ESCRT-0-III and the AAA-ATPase VPS4. Within this complex, ESCRT-III (named CHMPs in animal cells) and VPS4 manifest the minimal unit required for driving membrane fission (*2, 3*). ESCRTs have been shown to execute membrane fission in numerous cellular processes in eukaryotes, including cell division, multivesicular bodies formation (MVBs), vesicle release, membrane repair, neural development, and nuclear envelop resealing (*2, 4*). Additional non-canonical functions have also been proposed for this seminal complex (*5–9*). Although the involvement of ESCRTs in some of these processes has been documented in primitive cellular life forms (i.e. cell division and vesicle release), the original role that the ESCRT complex initially mediated is still elusive (*1, 10*).

Clues for the functional evolutionary origins of these proteins could reside in the prokarya domain that houses the most ancient life forms of our planet. In bacteria, distant ESCRT homologs have recently been identified, but their cellular function is still unknown (*11, 12*), while in archaea, the complex was associated with cell division and vesicle release (*1, 2, 8, 10, 13–17*). Recently, genes encoding for ESCRT proteins have also been found in Asgard archaea, a newly discovered Asgard super-phyla, that encode for several eukaryotic signature proteins (ESPs) and is currently considered as the ancestor of the first eukaryotic cell (*18–20*). To date, seven different ESCRT-related genes have been identified in Asgard archaea: homologs of two ESCRT-I proteins, two ESCRT-IIs, two ESCRT-IIIs (referred to here as CHMP 1-3, CHMP 4-7), and VPS4 (*18, 20*). While homologs for ESCRT-III and VPS4 were previously identified in archaea of the TACK superphyla (named Cdv B and Cdv C, respectively) (*1, 14, 15, 21*), several lines of evidence suggest that Asgard ESCRTs are more closely related to eukaryotes. First, homologs for ESCRT-I and –II have only been found in Asgard archaea. Second, domain analysis indicates that some domains of the Asgard ESCRTs are conserved in eukaryotes but are absent in other archaeal phyla (*22, 23*). Finally, in recent experiments performed in yeast the Asgard VPS4 homolog was significantly more efficient than the other archaeal homologs in reverting phenotypes associated with VPS4 absence (*22*). Hence, elucidating the function of Asgard ESCRTs can shed light on the ancient cellular function of this conserved membrane remodeling machine in eukaryotes.

The Asgard super-phyla is highly diverse and encompasses a growing number of phyla, including Lokiarchaeota (Loki), Thorarchaeota (Thor), Odinarchaeota (Odi), Helarchaeota (Hela), Freyarcheota (Frey), and Heimdallarchaeota (Heimdall) (*24*). Among these phyla, Loki was the first to be identified and is considered to be the most abundant, and Heimdall is thought to be the closest relative to eukaryotes *(18, 20)*. So far, most of the knowledge obtained on Asgard archaea was derived from metagenome sequence analysis. Moreover, there are currently no tools or appropriate model systems to study the biology of the Asgard archaea super phylum with the exception of one recent enrichment culture of Lokiarchaeota species population (*25*). Hence, studying the functional role of ESCRT proteins encoded by Asgard archaea in their native environment is extremely challenging.

To overcome these challenges and to track back the functional origins of these cardinal proteins, we set to characterize Asgard ESCRTs via their heterologous expression in mammalian cells. In this study, we find evidence to suggest that the functional origins of these proteins involve DNA binding. We achieved the expression of the genes encoding the two Asgard ESCRT-III subfamilies (CHMP1-3 and CHMP4-7) and VPS4 from both Heimdall and Loki origin in mammalian tissue culture cells. We further show that these proteins are working in concert. ESCRTIII/VPS4 homologs from Lokiarchaeota interacted with one another within mammalian cells and changed their subcellular localization upon co-expression with their counterparts. Unexpectedly, co-expression of Loki CHMP 4-7 and VPS4 led to accumulation of these proteins in discrete foci in the nucleus. Consistently, CHMP 4-7 was found in the chromatin fraction and recruited VPS4 to chromatin upon co-expression. Notably, mammalian ESCRT-III/VPS4 proteins (particularly CHMP1) were also found in the chromatin fraction, with human CHMP1B organizing in nuclear foci and recruiting both human and Loki VPS4 to these foci when overexpressed in cells. Moreover, removing the homologous N terminal domains in both human and Asgard ESSRT-IIIs abolished their ability to organize in nuclear foci. Hence, human and Asgard ESCRTs associate with chromatin and functionally interact with one another inside mammalian cells, corroborating their evolutionary conserved functionality of these proteins.

Finally, recombinant Loki CHMP 4-7 and human CHMP1B were able to directly bind DNA in vitro, indicating direct binding of ESCRT-III to DNA of both Asgard and human ESCRT-IIIs. Collectively, our findings highlight an ancient evolutionary conserved role of the ESCRT proteins to work in concert to interact with chromatin.

## RESULTS

According to current models, the ESCRT-III and VPS4 complexes constitute the core components of the ESCRT machine that execute membrane fission (*26–29*). They do so by assembling ESCRT-III monomers into helical filaments and remodeling these filaments by VPS4. To track the functional eukaryotic origins of ESCRT protein family in the Asgard super-phyla, we first performed hierarchical clustering based on sequence homology and maximum likelihood phylogenetic analysis of these proteins together with their human homologs. Our analysis of Asgard and human ESCRT-III/VPS4 as deposited in the databases show that these genes share sequence homology clustering together with their human homologs into three subgroups according to the standard ESCRT protein classification (Fig. 1). One group is composed of VPS4 homologs, and two groups are composed of ESCRT-III proteins, as previously shown using smaller cohorts (*18, 22*). The twelve human ESCRT-IIIs were divided between these two ESCRT-III subgroups so that CHMPs 1, 2, 3, and IST1 resided in one cluster (named CHMP1-3) and CHMPs 4, 5, 6, 7 in the second (named CHMP 4-7). Both the maximum likelihood tree and the sequence identity clustering analyses (Fig. 1, A and B respectively) show that the VPS4 cluster confers the highest sequence identity and evolutionary conservation between species while the sequences in the two ESCRT-III clusters are more diverged, with the CHMP4-7 cluster exhibiting divergent clade of sequences annotated as Lokiarchaeota and Heimdallarchaeota. Therefore, while VPS4 is relatively evolutionary conserved between different archaea species, ESCRT-IIIs are highly diverse, querying a functional similarity for these homologs within the archaea phylum and between archaea and humans.

**Fig. 1.**
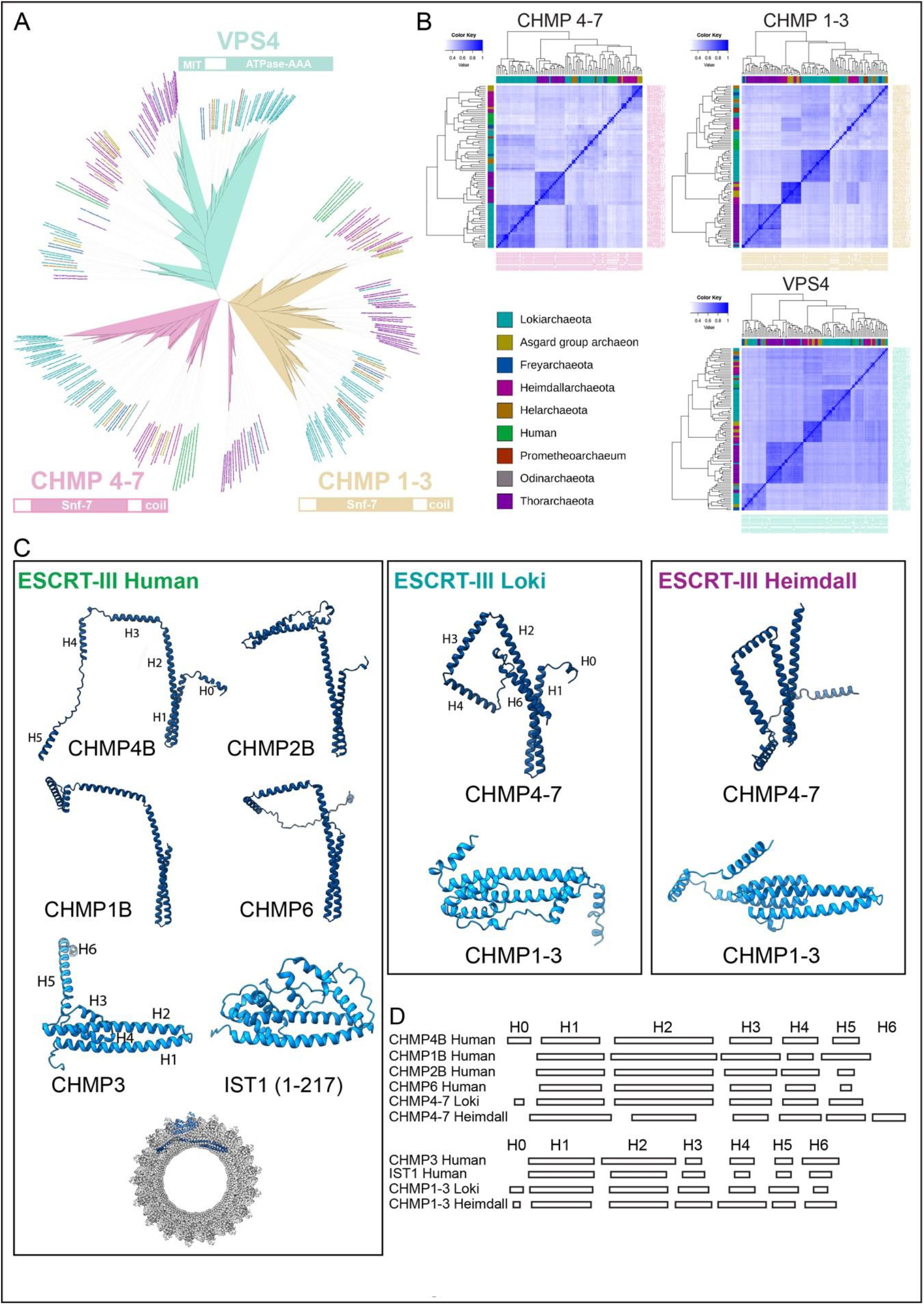
Phylogenetic and structural analysis of human and Asgard ESCRTIII/VPS4 (**A**) Maximum likelihood tree of human and Asgard ESCRT-III/VPS4. Colors of the branches denote the ESCRT classifications. Predicted domains for proteins in each cluster (InterPro) are denoted below cluster name. Leaf colors denote the phylogenetic associations of human and Asgard proteins (as indicated in colored index). (B) Heatmaps showing the sequence similarity of the three ESCRT clusters according to their classification on the phylogenetic tree, using Fitch–Margoliash distance. (**C)**) Selected Robetta predicted structures of human ESCRT-III proteins and the Asgard proteins examined (for confidence score, see fig. S1A). Full-length proteins are shown except for human IST1 in which only amino acids 1-217 are shown. Colors encode to “open” (dark blue) and “closed” (light blue) ESCRT-III conformations. Cryo-EM structure of the human IST1 and CHMP1B co-polymer (PDB ID, 3JC1) is shown below the human predictions. Single monomers of IST1, “closed” conformation (dark blue) and CHMP1B, “open” (light blue) are highlighted in the filament structure. Overlay of structure predictions of all human ESCRT-III proteins and the selected Asgard proteins from both Alphafold and Robetta as well as with the available structure in the protein database (PDB) is provided in fig. S1B. (**D**) Linear representation of the predicted helixes. Top and bottom panels, “open” and “closed” conformations, respectively. Helixes were numbered as illustrated for human ESCRT-IIIs (**C**, left panel). The length of each rectangle corresponds to the helix length.

Asgard ESCRT-III homologs from both clusters (CHMP1-3, CHMP4-7) were found to include the conserved ESCRT snf-7 polymerization domain and a C-terminal coil domain in domain analysis (Fig. 1A) and were reported to possess the conserved ESCRT protein-protein interaction motifs (MIM domains) (*22, 23*). Asgard VPS4s had the conserved MIT domain, which governs the interaction between VPS4 and ESCRT-III in eukaryotes (Fig. 1A) (*22, 30*). Therefore, suggesting that Asgard ESCRT-III homologs retain polymerization and interaction with VPS4.

Following this notion, we examined whether the overall structure of ESCRT-III Asgard is similar to their human homologs. For this purpose, we generated structural models for human ESCRT-III proteins and ESCRT-III homologs encoded by Loki and Heimdall species using the deep-learning based structure prediction servers Robetta and Alpha-fold (Fig. 1C and fig. S1) (*31, 32*). The 12 human ESCRT-III proteins clustered into two main folds, previously termed the “open” and “closed” conformations (*3*), with 10 ESCRT-III proteins obtaining the “open” conformation and two (CHMP3 and IST1) obtaining the “closed” conformation. Interestingly, the structural models obtained for Asgard ESCRT-III homologs of both Loki and Heimdall resembled those obtained for human ESCRT-IIIs with CHMP4-7 obtaining the so-called “open” conformation and CHMP1-3 the “closed” conformation. The number and length of helixes in proteins that exhibited similar conformation (“open” or “closed”) were also similar between human and Asgard ESCRT-III homologs, except for Heimdall CHMP4-7, which had shorter helixes and an additional helix at the C terminal (Fig. 1D). Hence, human and Asgard ESCRT-III homologs appear to share similar 3D structural folds despite the low sequence homology and the evolutionary gap that separate them, supporting polymerization capabilities for Asgard ESCRT-IIIs. Moreover, the “open” and “closed” conformations of ESCRT-III proteins appear to be evolutionary conserved and may therefore be crucial for the function of the machinery.

As our predictions show that the human and Asgard ESCRT-III homologs share structural similarities, we set forward to explore their function in mammalian and human tissue culture systems. To this end, we sub-cloned genes encoding Loki and Heimdall ESCRT-III proteins (CHMP1-3, CHMP4-7) and VPS4 into expression vectors (three proteins from each family). We specifically chose ESCRT-III and VPS4 genes proteins that are coded from the same metagenomic assembled genomes (MAGs) to increase the likelihood of ESCRTs to interact with one another and carry out their function.

We successfully expressed all six ESCRT proteins in mammalian and human tissue culture cells (Fig. 2A). We find that ESCRT-III and VPS4 proteins are situated in different localities within the cell and that the cellular distribution of Loki and Heimdall ESCRTs is slightly different. Loki ESCRTs were found in both the soluble and insoluble fraction with CHMP1-3 and VPS4 enriched in the soluble fraction and CHMP4-7 equally distributed in both fractions. Heimdall ESCRTs CHMP4-7 and VPS4 were predominantly in the soluble fraction, while CHMP1-3 was found to be enriched in the insoluble fraction (Fig. 2B). In confocal images, fluorescently tagged versions of Heimdall ESCRTs were found to reside in both the nucleus and cytosol. Loki VPS4 was also distributed in both compartments but Loki CHMP1-3 was mainly cytosolic and Loki CHMP4-7 was predominantly nuclear (Fig. 2C). These results were observed in several cultured cell lines indicating that they reflect the general tendency of the proteins to distribute inside cells (fig. S2A). Therefore, genes encoding Asgard ESCRTs are expressed in human and mammalian cells and reside in both the nucleus and the cytoplasm. While these findings are consistent with the ability of human ESCRT-III/VPS4 to distribute between the cytosol and nucleus, the realization that ESCRT homologs encoded by prokaryotic Asgard archaea enter the eukaryotic nucleus, is noteworthy.

**Fig. 2.**
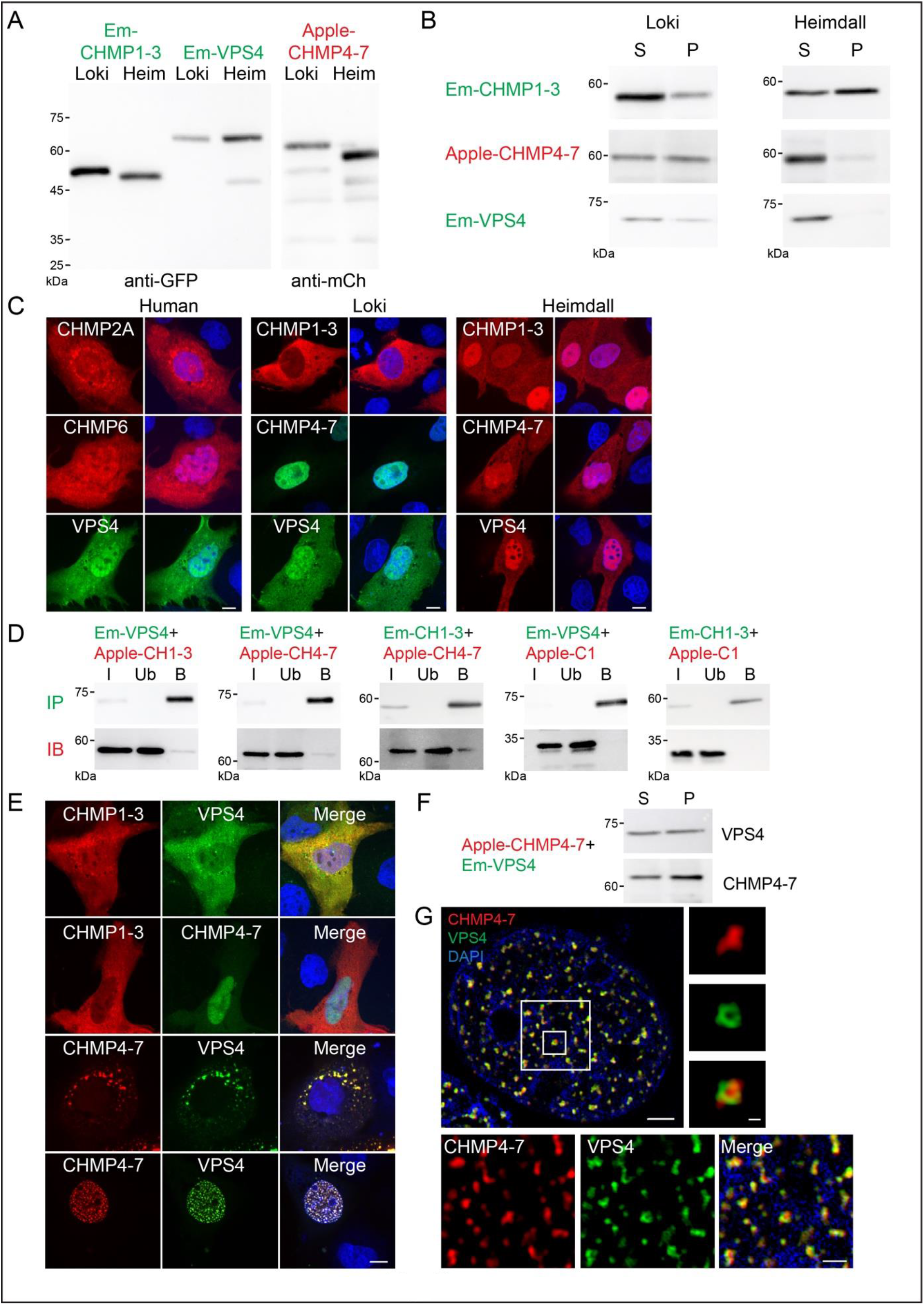
ESCRTIII/VPS4 homologs from Asgard archaea express and interact with one another inside mammalian cells. MDCK cells were transfected with Emerald-CHMP1-3, Apple-CHMP4-7, or Emerald-VPS4 homologs from either Loki or Heimdall. (**A-B**) cells were lysed (A) or lysed and fractionated to supernatant (S) and pellet (P) (**B**), and subjected to western blot analysis using anti-mCh or anti-GFP antibodies. (**C**) Fluorescently tagged versions of ESCRTIII/VPS4 proteins from Human (left), Loki (middle), and Heimdall (right) were expressed in MDCK cells, as indicated. Followed fixation and Hoechst staining (shown in merged images) cells were imaged using confocal microscopy. Note that both Loki and Heimdall ESCRT homologs can enter the eukaryotic nucleus. Representative single slice images are shown. Data obtained from at least three independent experiments, n≥ 50 for each condition, scale = 10 μM. Similar results were obtained in HeLa and NIH3T3 cells (fig. S2A). (**D**) MDCK cells were co-transfected with fluorescently tagged Loki ESCRTIII/VPS4 combinations, as indicated, subjected to immunoprecipitation using anti-GFP antibodies, and immunoblotted (IB) using anti-mCherry antibodies. (I- cell extracts, Ub -unbound fraction, B- bound fraction). Data was reproduced in at least two independent experiments. (**E**) MDCK cells were co-transfected with fluorescently tagged Loki ESCRTIII/VPS4, as indicated. Followed fixation and Hoechst staining (shown in merged images), cells were imaged using confocal microscopy. Scale = 10 μM. Representative single slice images are shown. Note that co-expression of CHMP4-7 and VPS4 induced the accumulation of both proteins at discrete foci in the nucleus (~ 30% of the cells). Images were obtained from at least three independent experiments, n≥ 100 for each condition. Similar results were obtained in HeLa and NIH3T3 cells (fig. S2D). (**F**) Sedimentation analysis of MDCK co-transfected with Loki Apple CHMP4-7 and Emerald VPS4. Both VPS4 and CHMP4-7 translocated to the pellet fraction. Data were reproduced in at least two independent experiments. (**G**) SIM images of MDCK cells co-transfected with Loki Apple-CHMP4-7 and Emerald-VPS4 and stained with DAPI. Representative single slice images are shown. White boxes indicated zoomed-in images. Large box, bottom panel; small box, right panel. Scale = 2 μM (top left panel), 1 μM (bottom panel), 0.2 μM (right panel). n=10.

To find out whether Asgard ESCRTs can interact with one another in the cellular milieu, we co-expressed pairs of ESCRT-III/VPS4 proteins in mammalian cells. Co-IP experiments revealed that all three ESCRT Loki homologs bind to one another inside cells (Fig. 2D). However, no interaction between Heimdall ESCRT homologs was detected (fig. S2B). Therefore, while ESCRT homologs from both Loki and Heimdall are expressed in mammalian cells, only the Loki homologs preserved their ability to interact with one another in mammalian cells.

Given the ability of Loki ESCRT-III/VPS4 proteins to interact, we examined the subcellular localization of these proteins upon co-expression. In our experimental setup, most co-expression conditions did not affect the subcellular distribution documented for each protein when expressed alone (Fig. 2E). Co-expression of CHMP 4-7 and VPS4 homologs from Loki, but not Heimdall, induced co-localization of CHMP 4-7 and VPS4 in discrete puncta in the cytosol, resembling the localization pattern observed for mammalian ESCRTs in multi vesicular bodies (Fig. 2E and fig. S2C) (*33*). Consistently, under these conditions, CHMP4-7 was shifted to the Pellet fraction, indicating its increased association with cellular membranes (Fig. 2F). Strikingly, prominent co-localization between Loki CHMP4-7 and VPS4 was also observed in the nucleus, with both proteins accumulating in discrete puncta decorating the entire nucleus (~30% of the co-transfected cells). In structured illumination microscopy (SIM) images, VPS4 and CHMP4-7 appeared to co-reside in nuclear puncta with VPS4 engulfing CHMP4-7, confirming co-localization between the proteins and suggesting an ordered organization of the complex inside the nucleus (Fig. 2G). This phenotype was not reproduced upon co-expression of Heimdall CHMP4-7 and VPS4 but was reproduced for Loki ESCRTs in MDCK, NIH3T3, and HeLa cells (Fig. 2E and fig. S2, C and D), suggesting that it reflects a phenotype that is specific to Loki ESCRTs. Moreover, CHMP4-7 and VPS4 proteins interacted with one another to facilitate the formation of nuclear foci only in an interspecies manner, as nuclear foci were not observed upon co-expression of Loki CHMP4-7 with Heimdall VPS4 or vice versa (fig. S2E). Together with the different cellular distributions observed for these homologs (Fig. 2C), our results suggest that Heimdall and Loki ESCRTs exhibit distinct behaviors within cells and exclusive interspecies interaction in mammalian cells.

Intrigued by the striking co-localization of Loki CHMP4-7 and VPS4 to discrete foci in the nucleus, we set to examine the ability of Asgard ESCRT-III/VPS4 proteins to interact with one another and associate with chromatin using a fractionation assay. We find that both Loki CHMP4-7 and CHMP1-3 reside in the chromatin-bound fraction either when expressed alone or in the presence of an additional ESCRT partner. Loki VPS4, on the other hand, was found at low levels in the chromatin fraction, when expressed alone or together with CHMP1-3, but exhibited prominent chromatin association in the presence of CHMP4-7 (Fig. 3A). Heimdall CHMP4-7 and VPS4 were found in negligible amounts in the chromatin fraction, explaining the lack of ESCRT-III/VPS4 nuclear foci in cells (fig. S2F). Therefore, Loki CHMP4-7 resides in the chromatin fraction and recruit Loki VPS4 to chromatin upon their co-expression (a model for the interplay of Loki ESCRT-III/VPS4 in mammalian cells is presented in Fig. 2B).

**Fig. 3.**
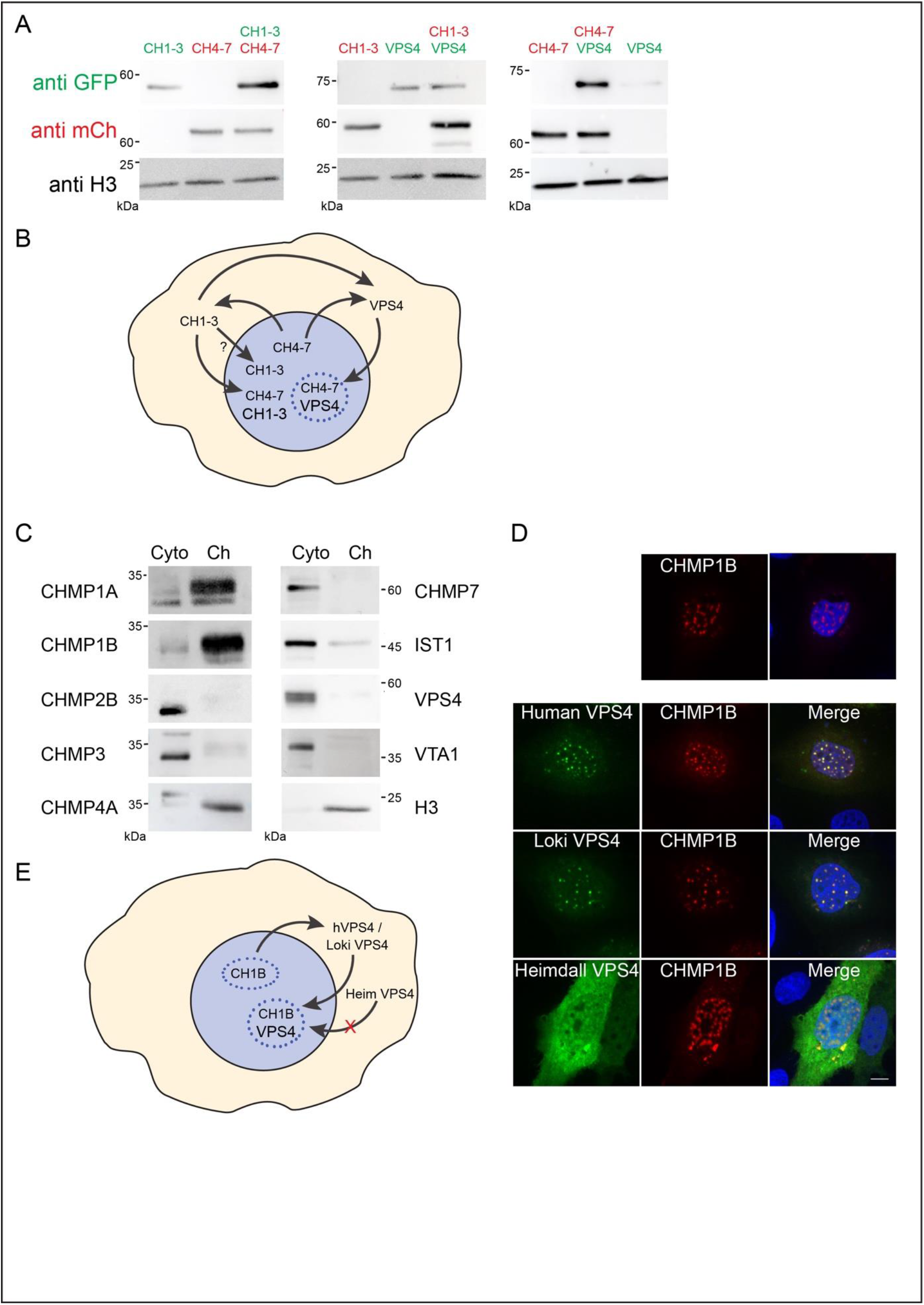
Loki and human ESCRT-III/VPS4 proteins associate with chromatin. (**A**) MDCK cells were transfected or co-transfected with fluorescently tagged versions of ESCRT-III/VPS4 Loki homologs (Emerald-CHMP1-3; CH1-3, Apple-CHMP4-7; CH4-7, Apple-CHMP1-3; CH1-3 or Emerald-VPS4). Cells were then fractionated using protein fractionation Kit (see Methods) and normalized portions of chromatin extracts (20 μg) were loaded and analyzed by western blot. H3 (histone 3) staining was used as chromatin-bound marker. Data were reproduced in at least three independent experiments. Top middle panel was subjected to longer exposure times relative to other panels due to extremely low levels of VPS4 in the chromatin fraction. (**B**) Schematic illustration of the interplay between Loki ESCRTIII/VPS4 proteins. All three proteins can interact with one another. CHMP4-7 and CHMP1-3 associate with chromatin, while VPS4 does not. Co-expression of CHMP4-7 with either CHMP1-3 or VPS4, increase their association with chromatin (indicated by the larger font size), but does not affect CHMP4-7 levels in chromatin. Only co-expression of CHMP4-7 and VPS4 results accumulation in nuclear foci (indicated by blue dots). (**C**) Naïve MDCK cells were fractionated, and normalized amounts of cytoplasm (Cyto) and chromatin (Ch) extracts (20 μg) were loaded and subjected to western blot analysis using specific antibodies for the indicated proteins (see table S1). Data were reproduced in at least two independent experiments. (**D**) Expression of human HA-CHMP1B alone (upper panner) or together with Emerald-VPS4 human, Loki or Heimdall proteins (lower panel) in MDCK cells. Cells were stained with anti-HA antibodies and imaged using confocal microscopy. Representative single slice images are shown. Scale = 10 μM. Images were obtained from at least two independent experiments, n≥ 30 for each condition. (**E**) Schematic illustration summarizing the interactions between human CHMP1B and different VPS4 homologs. CHMP1B accumulates at discrete foci in the nucleus and recruit human- and Loki-, but not Heimdall-VPS4 to these foci.

Few endogenous ESCRT-III proteins, particularly CHMP1A and CHMP1B, were also found to reside predominantly in the chromatin fraction (Fig. 3C). Moreover, overexpression of CHMP1B gave rise to discrete foci in the nucleus with human VPS4 recruited to these Foci (Fig. 3D).

Strikingly, Loki VPS4, but not Heimdall VPS4, was also recruited to CHMP1B nuclear foci upon its co-expression with human CHMP1B (Fig. 3, D and E). We, therefore, concluded that: (a) the chromatin binding capabilities of ESCRT-III is conserved in humans and Asgard archaea and (b) human CHMP1B is capable of recruiting Loki VPS4 to nuclear foci, suggesting that human ESCRT-IIIs can interact with some of their Asgard VPS4 despite more than a billion years of evolution.

Next, we aimed to elucidate the basis for Loki ESCRT-III/VPS4 chromatin association and recruitment (Fig. 4, A to I). The MIT domain in VPS4 is known to mediate VPS4 interaction with ESCRT-III in eukaryotes and was found to be conserved in Asgard VPS4 homologs (Fig. 1A) (*22, 30*). We, therefore, asked how depleting the MIT domain in VPS4 will affect the ability of CHMP4-7 to recruit VPS4 to chromatin and form nuclear foci. MIT-depleted fluorescently tagged Loki-VPS4, expressed in cells and distributed similarly to full-length Loki-VPS4 (Fig. 4A and fig. S3A). In chromatin fractionation assay, Loki VPS4-ΔMIT was still recruited to chromatin in the presence of CHMP 4-7, suggesting that the MIT domain is dispensable for the recruitment of VPS4 to chromatin (Fig. 4B). In confocal images, however, VPS4-ΔMIT co-localized with CHMP4-7 in discrete foci outside the nucleus, but the percentage of cells exhibiting VPS4/CHMP4-7 puncta in the nucleus was considerably reduced (Fig. 4, C and F). Loki VPS4-ΔMIT was not recruited to CHMP1B nuclear foci upon their co-expression, indicating that Loki VPS4 interacts with human CHMP1B via its MIT domain (fig. S3B). These data suggest that the MIT domain in Loki VPS4 is conserved enough to interact with human ESCRT-III proteins and is essential for the induction of nuclear foci by Loki ESCRT-III/VPS4 proteins. Yet, additional regions in Loki VPS4 contribute to its interaction with Loki CHMP 4-7, allowing it to interact with- and be recruited by-Loki CHMP4-7 in the absence of the MIT domain.

**Fig. 4.**
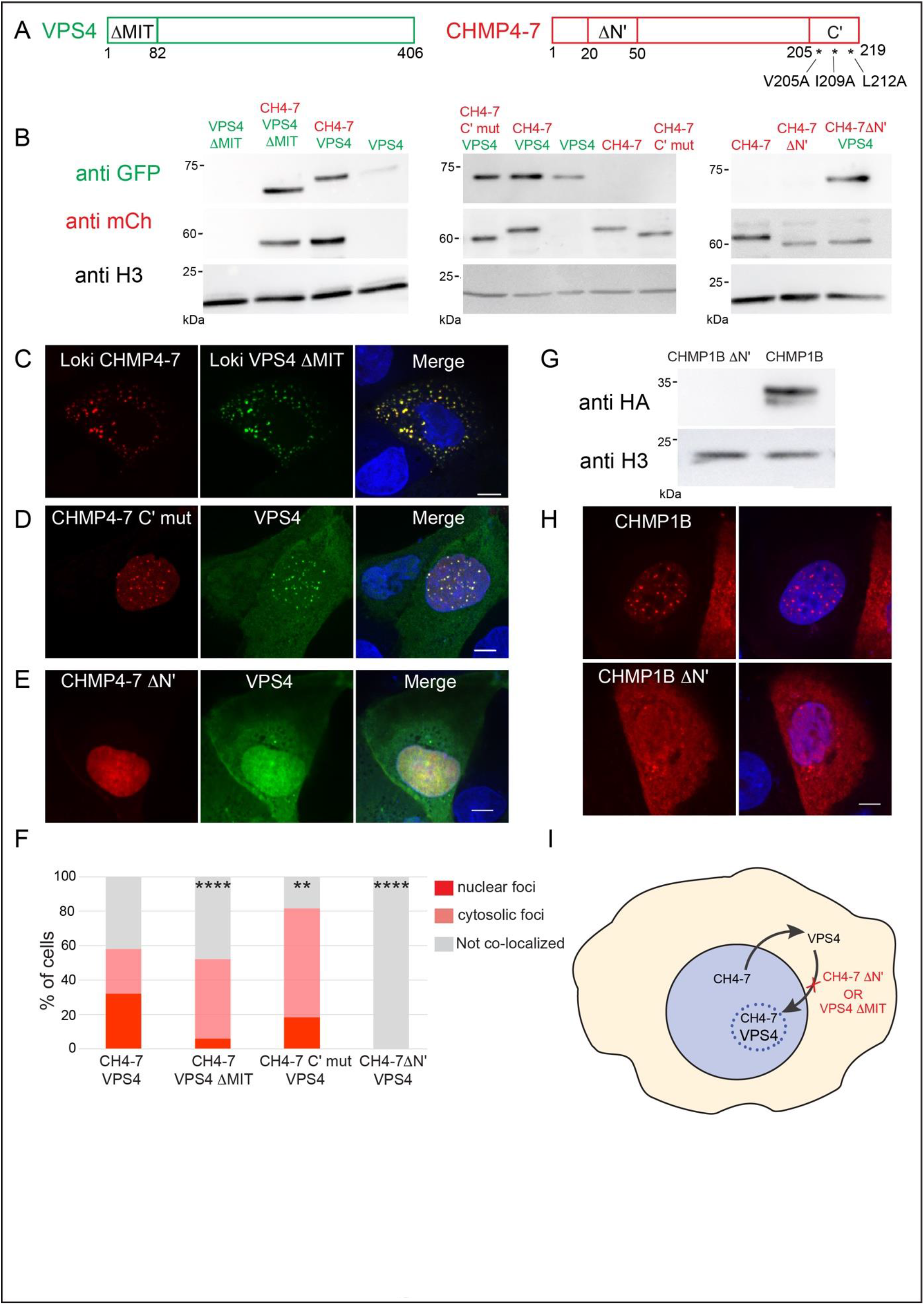
The N-terminal region in ESCRT-III mediates chromatin association and nuclear foci formation in both human and Loki homologs. (**A**) Schematic illustration of VPS4 and CHMP4-7 mutants used in panels B-I. The indicated regions were either deleted or replaced, as shown. (**B**) Chromatin fractionation. MDCK cells transfected with the indicated fluorescently tagged ESCRT-III/VPS4 Loki mutants were fractionated, and normalized amounts of chromatin extracts (20 μg) were loaded and analyzed by western blot. Data were obtained from two independent experiments. (**C-E**) Representative single-slice images of cells transfected with fluorescently tagged versions of the indicated ESCRT-III/VPS4 mutants, imaged using confocal microscopy. Scale = 10 μM. Data were obtained from at least three independent experiments, n≥ 60 for each condition. (**F**) Percentages of cells that exhibited foci in the nucleus, co-localization in the cytoplasm or no co-localization, upon co-transfection with the indicated plasmids (Apple-CHMP4-7 and Emerald-VPS4, n=73, Apple-CHMP4-7 and Emerald-VPS4ΔMIT, n=60, Apple-CHMP4-7 NES mut and Emerald-VPS4, n=75, Apple-CHMP4-7 ΔN’ and Emerald-VPS4, n=73). Note that deleting the MIT domain in VPS4 reduced accumulation in nuclear foci, while deleting the N’ domain in CHMP4-7 completely abolished both the interaction between the two proteins and the formation of nuclear foci. Data were obtained from at least three independent experiments. Statistical analysis of the foci in the nucleus was calculated using Chi-square test two tail (**p- value ≤ 0.01, ****p- value ≤ 0.0001). (**G-H**) MDCK cells were transfected with human HA-CHMP1B or with CHMP1B N’ mutant in which the homologue residues of Loki CHMP4-7 were deleted (CHMP1B ΔN). Cells were subjected to chromatin fractionation assay (**G**) or imaged using confocal microscopy (**F**). (**G**) Cells were fractionated, and normalized portions of chromatin extracts (20μg) were analyzed by western blotting using HA antibodies. Data were reproduced in at least two independent experiments. (**H**) Representative single slice images of cells imaged by confocal microscopy. Scale = 10 μM. Images obtained from at least two independent experiments, n≥ 40 for each condition. Note that CHMP1B ΔN mutant did not reside in the chromatin fraction and did not accumulate in nuclear foci. (**I**) Schematic illustration of the interactions between Loki VPS4 and CHMP4-7 mutants. Co-expression of CHMP4-7 and VPS4 induced accumulation of these proteins in nuclear foci, deletion of either N’ from CHMP4-7 or MIT domain from VPS4 abolish this phenotype.

To identify putative chromatin binding regions in ESCRT-III, we performed multiple sequence alignment between Loki CHMP4-7 and human CHMP1 isoforms (fig. S3C). This comparison highlighted conserved regions in the N and C terminals of the proteins. Introducing point mutations to C-terminal in CHMP4-7 did not affect the ability of CHMP4-7 to associate with chromatin nor its ability to recruit VPS4 to chromatin, suggesting that the C-terminal of CHMP4-7 is not involved in chromatin binding nor in recruiting VPS4 to chromatin (Fig. 4, A and B). Consistently, CHMP4-7/VPS4 nuclear foci could be readily detected in cells co-expressing CHMP4-7 C’ mutants together with VPS4 (Fig. 4, D and F, ~20% in mutant compared to ~30 in WT). Depleting the N-terminal, however, resulted in reduced association of CHMP4-7 with chromatin and a complete loss of CHMP4-7/VPS4 nuclear foci (Fig. 4, A, B, E and F), but did not affect the nuclear localization of CHMP4-7 (fig. S3A). These results indicate that the N-terminal region in CHMP4-7 is specifically involved in its ability to associate with chromatin in cells. Depleting the parallel region in human CHMP1B also abolished its ability to associate with chromatin and to organize in discrete nuclear foci (Fig. 4, G and H), indicating that N terminal domain identified here governs chromatin association in both human and Loki ESCRT-III proteins (a summary of the interactions between Loki CHMP4-7 and VPS4 mutants is presented in Fig. 4I).

Finally, we set to examine the ability of ESCRT-III proteins to bind DNA directly. For that, we expressed and purified Loki CHMP4-7 and human CHMP1B from *E. coli* and performed a gel shift assay using a 5’ Cy5-tagged 40 nucleotides double-strand DNA. Under these conditions, both Loki CHMP4-7 and human CHMP1B were found to bind DNA, with CHMP1B showing stronger association including at lower protein concentrations (Fig. 5A). These results strongly suggest that the evolutionary conserved chromatin association observed here ESCRT-III is driven by direct binding to DNA.

**Fig. 5.**
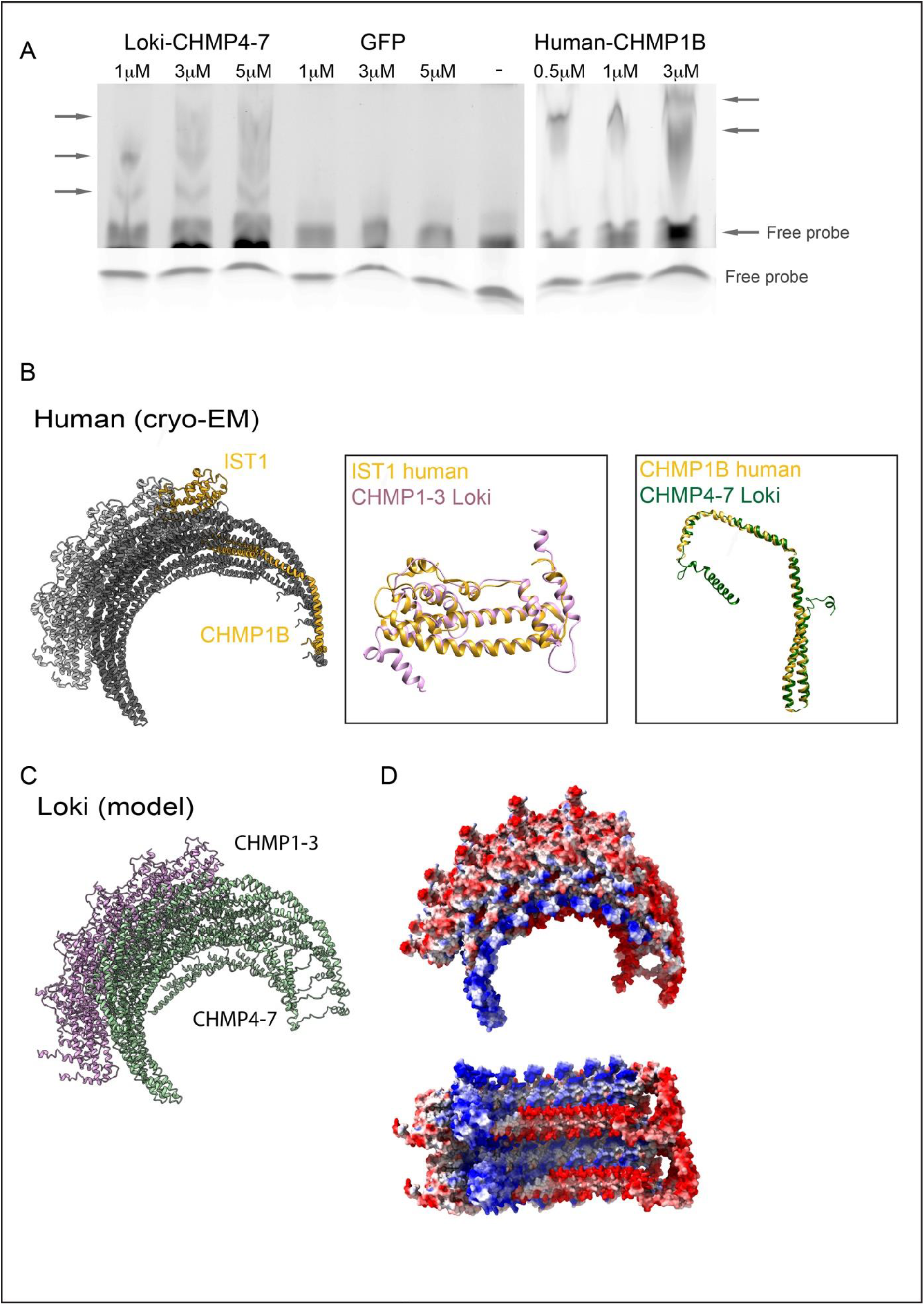
Loki CHMP4-7 and human CHMP1B directly bind DNA. (**A**) Electrophoretic mobility shift assay (EMSA) performed in the presence of Cy5-5’ end labelled double-stranded DNA (2.5 μM, 40bp) and the purified proteins: Loki CHMP4-7 (1, 3, 5 μM), GFP (negative control; 1, 3, 5 μM), or human CHMP1B (positive control, 0.5, 1, 3, μM). (−) = no protein. Arrows in upper panel indicate complex formation. Lower panel, free DNA probe as seen by shorter exposure time. Data were reproduced in at least three independent experiments. (**B-D**) Predicted structural organization of Loki ESCRT-III filament. (**B**) Left: Sectioned representation of the cryo-EM human IST1 and CHMP1B co-polymer structure (PDB ID 3JC1). Middle to right panels: overlay of the predicted structure of Loki CHMP 1-3 (Robbeta) on the human IST1 cryo-EM monomer structure, and the predicted Loki CHMP 4-7 structure (Robbeta) on human CHMP1B cryo-EM monomer structure, respectively. (**C**) The predicted Loki CHMP 1-3 and CHMP 4-7 structures (Robetta) were overlaid on the co-polymer cryo-EM structure of human IST1 and CHMP1B, respectively, to generate a theoretical 3D structural organization model of Loki ESCRT-III proteins. Shown are Loki CHMP1-3 (purple) and Loki CHMP 4-7 (green). (**D**) Electrostatic map of the theoretical Loki ESCRT-III co-polymer. Positive charge, blue; Negative charge, red; Neutral, white. Top panel, side view; Bottom panel axial view, showing the tube from the inside. Note, the accumulation of positively charged amino acids in the inner side of the tube which may infer affinity to negatively charged DNA.

## DISCUSSION

We propose that ESCRT-III proteins exhibit evolutionary conservation of chromatin / DNA binding properties between Asgard archaea and human lineages and provide evidence for the involvement of conserved protein-protein interactions in this process.

Our study was based on the heterogeneous expression of predicted coding sequences from Asgard archaea in mammalian cells. We found that genes encoding Asgard ESCRT-III and VPS4 homologs are successfully expressed in several cell lines and exhibit similar subcellular distributions. Moreover, Loki ESCRT-III/VPS4 proteins were found to interact with one another inside mammalian cells. These results indicate that Asgard ESCRT proteins can be studied inside mammalian ESCRTs and suggest that heterologous expression of Asgard genes in mammalian cells can be used as a tool to study the biochemical and cellular properties of ESPs encoded by Asgard archaea.

The low sequence identity values observed between predicted ESCRT-III proteins from different Asgard species (Fig. 1, A and B) motivated us to sample ESCRT-III/VPS4 modules taken from two different species. At the same time, as eukaryotic ESCRTs rely on an intimate protein-protein interactions network to mediate their function, we specifically chose coding sequences obtained from the same MAG. Indeed, we found that ESCRT-III/VPS4 Loki and Heimdall homologs exhibited different cellular properties and did not interact with one another inside cells. Overall, Loki ESCRT-III proteins appeared to be more functional than Heimdall in mammalian cells. Additionally, Loki-VPS4 was recruited to CHMP1B nuclear foci, while Heimdall VPS4 failed to do so. Hence, although based on phylogeny, Heimdall archaea are considered closer to humans, our results suggest that Loki ESCRTs may be functionally closer. Interestingly, while the predicted structure of Loki CHMP4-7 highly resembled the overall fold of the human ESCRT-III “open” conformation, Heimdall CHMP 4-7 had a slightly different fold, exhibiting shorter helixes and an additional C-terminal helix (Fig. 1, C and D). It is, therefore, possible that the basis for the conservation of function in the ESCRT kingdom lye in their structural similarity more than in their sequence identity.

The predictions obtained by the deep-learning based servers Alpha-Fold and Robetta highlighted conserved structural features of ESCRT-III proteins. Traditionally, ESCRT-III proteins were shown to obtain an autoinhibitory loop that opens upon activation and allows polymerization of the ESCRT-III filament (*3, 33*). The inactive and active conformations were termed “closed” and “open”, respectively. Among the twelve human ESCRT-III proteins, ten were predicted to have an “open” conformation and two a “closed” conformation. Interestingly, in both Asgard species used in this study, one ESCRT-III protein was predicted to be in an “open” conformation and one in “closed”, suggesting that both conformations are conserved in Asgard ESCRT-IIIs (Fig. 1, C and D and fig. S1). Interestingly, the structures of eukaryotic co-polymers described so far were composed of ESCRT-III pairs exhibiting “open” and “closed” conformations (CHMP2A and CHMP3, CHMP1B and IST1, respectively) (*34, 35*), raising the possibility that both conformations are needed for building a functional ESCRT filament. To test this notion, we overlaid the deep-learning based structures we generated for Loki CHMP1-3 and CHMP4-7 on the solved cryo-EM structure of the CHMP1B-IST1 co-polymer (PDB ID 3JC1, Fig. 5B) (*34*). Specifically, we overlaid Loki CHMP1-3 on IST1, both in “closed” conformation and Loki CHMP4-7 with CHMP1B, both in “open” conformation. Despite extremely low sequence identity values (20.74% between Loki CHMP1-3 and IST1 and 17.52% between Loki CHMP 4-7 and CHMP1B), a very good fit was observed between the Loki and human proteins, which enabled us to generate a putative structural model for Loki ESCRT-III filaments (Fig. 5C).

Mapping the electrostatic surface of the modeled Loki ESCRT-III filament, revealed a patch of positively charged amino acids that faces its inner surface (Fig. 5D). This positive patch of amino acids is part of the N-terminal helix of CHMP4-7, which we found to govern CHMP4-7-chromatin association. Moreover, this region is equivalent to the N-terminal helix in CHMP1B, which include a patch of positively charged amino acids that were suggested to drive its interactions with nucleic acids, *in vitro* (*36*). It is, therefore, tempting to speculate that, similar to their human homologs, Loki ESCRT-III organize into helical filament and are able to bind DNA through electrostatic interactions with positive amino acids that reside in the inner side of the filament. However, this notion currently relies on theoretical models, and further experimental work is required to substantiate it.

Our findings that ESCRT-IIIs associate with chromatin are in line with previous findings in mammalian cells. In 2001 Stauffer and colleagues reported that CHMP1 concentrates at chromatin-condensed regions and protects them from nuclease degradation (*5*). More recently, a truncated version of CHMP7 was shown to accumulate in discrete foci in the nucleus and to recruit CHMP4B to these foci (*37*), and a cryo-EM structure of IST1 and CHMP1B filament was solved in the presence of nucleic acid, showing direct interactions between the ESCRT-III filament and nucleic acids (*34*). Our findings that these properties are shared in both Asgard and mammalian ESCRTs and are mediated by the same domain in the human and Loki homologs strongly suggest that the ability of ESCRT-III proteins to bind DNA is physiologically relevant.

Our data point to the involvement of chromatin binding in the function of ancient ESCRTs. ESCRTs have been shown to be involved in cell division in both TACK archaea and in eukaryotes (*1, 10*). In TACK archaea, CdvA and the ESCRT-III-VPS4 homologs CdvB and CdvC have been designated as the cell division machinery (*14, 15, 21*), and CdvA was found to assemble into filaments in the presence of DNA (*38*). Given the established role of ESCRT in membrane remodeling (*4, 39*), it is possible that ESCRTs were originally evolved as a basic machinery for driving cell division by coupling DNA segregation and membrane constriction.

An alternative explanation to the observed data is that ESCRTs may have a conserved role in genome stability and activity. Eukaryotic ESCRT-IIIs have been documented to be involved in genome retrotransposition via the LINE complex and to localize to micronuclei (*6, 37, 40*). In addition, CHMP1B was shown to reside in chromatin-condensed regions and protect them from nuclease degradation (*5*), and CHMP2A (also called BC-2) was shown to accumulate in discrete regions in the nucleus where it co-localized with the histone3 phosphorylation marker PH3 and with the transcriptional silencing protein BM1 (*7*). Lastly, ESCRTs have been identified as nuclear matrix and chromatin binding proteins in several large-scale proteomic screens (*40–42*). Whether these high genome organization properties exist in Asgard archaea is still unknown. Nevertheless, our findings that Loki CHMP4-7 can enter the nucleus and recruit VPS4 to the nucleus in mammalians suggest that Asgard ESCRT-IIIs can enter the nuclear membrane. In this respect, Loki and Heimdall archaea enriched from sediments in Arhus Bay were recently reported to have a condensed DNA organization (*43*). Although a membrane engulfing these condensed DNA structures has not been shown yet, it is tempting to speculate that the ability of Asgard ESCRTs to enter the mammalian nuclear membrane that we have documented represent their physiological properties in archaea and that the evolutionarily conserved role of ESCRTs involves genome stability and protection.

## ACKNOWLEDGMENTS

We thank Dan Levi (BGU) and Eran Meshorer (HUJI) for fruitful discussions on chromatin binding proteins and constructive advice. We also thank, Dr. Ran Zalk (BGU) for helpful feedback and technical help with structural analysis. We thank all members of the Elia lab for critical feedback throughout the project. The Elia laboratory is funded by the Israeli Science Foundation (ISF) Grant no. 1323/18. I. M. acknowledges support from the European Research Council under the European Union’s Horizon 2020 research and innovation program (grant no. 640384) and the Israel Science Foundation (grant no. 1947/19).

## AUTHOR CONTRIBUTIONS

DN, IM and NE conceptualized the project. DN designed, performed and analyzed all cell biology and biochemical experiments. NM purified recombinant proteins. AZ performed phylogenetic analysis. YD assisted in experiments. RZ generated structural model and helped with structural data interpretation. NE wrote the manuscript with the help of DN and IM.

